# Contribution of Lateral Gene Transfer to the evolution of the eukaryotic fungus *Piromyces sp*. E2: Massive bacterial transfer of genes involved in carbohydrate metabolism

**DOI:** 10.1101/514042

**Authors:** Isabel Duarte, Martijn A. Huynen

## Abstract

Lateral gene transfer (also known as Horizontal Gene Transfer) is the transmission of genetic material between phylogenetically unrelated organisms. Previous studies have been showing the importance of this process for the evolution of unicellular eukaryotes, particularly those living in highly competitive niches such as the herbivore gut.

*Pyromices sp*. is an obligate anaerobic chytrid fungus that grows as a commensal organism in the gut of mammalian herbivores, possessing hydrogenosomes instead of mitochondria, producing hydrogen, and playing a key role in the digestion of plant cell wall material. These particular features make its genome particularly valuable for the study of the evolution and adaptation of unicellular eukaryotes to the cellulose-rich and anaerobic environment of the herbivore gut.

Here we performed a detailed large-scale lateral gene transfer (LGT) analysis of the genome from the chytrid fungus *Piromyces sp*. strain E2. For this we set out to elucidate (i) which proteins were likely transferred to its genome, (ii) from which bacterial donor species, and (iii) which functions were laterally acquired. Using sequence comparison and phylogenetic analyses, we have found 704 LGT candidates, representing nearly 5% of the *Piromyces sp*. orfeome (i.e. the complete set of open reading frames), mostly transferred from Firmicutes, Fibrobacteres, Bacteroidetes and Proteobacteria, closely following the microbial abundance reported for the herbivore gut. With respect to the functional analysis, the LGT candidate set includes proteins from 250 different orthologous groups, with a clear over-representation of genes belonging to the Carbohydrate Transport and Metabolism functional class. Finally, we performed a graph density analysis on the metabolic pathways formed by the LGT candidate proteins, showing that the acquired functions fit cohesively within Piromyces metabolic network, and are not randomly distributed within the global KEGG metabolic map. Overall, our study suggests that Piromyces’ adaptation to living anaerobically and in the a cellulose-rich environment has been undoubtedly fostered by the acquisition of foreign genes from bacterial neighbors, showing the global importance of such evolutionary mechanisms for successful eukaryotic thriving in such competitive environments.

## Introduction

Lateral gene transfer is the non-genealogical transmission of genetic material from one organism to another [1]. This sharing of genetic material between phylogenetically unrelated organisms fuels the fast acquisition of new functions, increasing the metabolic toolkit of the receiving organism. Accordingly, it is currently recognized as a major evolutionary force shaping a species’ genomic content, particularly in single celled organisms [2, 3]. While this process has been well documented for transfers between prokaryotic species [4, 5, 6, 7], the role of this process in eukaryotic evolution remains controversial [8, 9], especially when multicellular eukaryotes, and humans in particular, are concerned.

In fact, the first draft of the human genome, in 2001, reports a set of 223 proteins (113 confirmed by PCR not to be contaminations) “that have significant similarity to proteins from bacteria, but no comparable similarity to proteins from yeast, worm, fly and mustard weed, or indeed from any other (nonvertebrate) eukaryote” [10]. This report immediately prompted two response papers [11, 12] questioning those results (which were classified based on best BLAST hits). Salzberg et al., (2001) provided a re-analysis of the data, where the genes from the initial set were systematically excluded with each step of the re-analysis, ending up with a group of circa 40 candidate horizontal gene transfers. Despite the largely reduced set, Salzberg et al. conclude by stating: “The more probable explanation for the existence of genes shared by humans and prokaryotes, but missing in nonvertebrates, is a combination of evolutionary rate variation, the small sample of nonvertebrate genomes, and gene loss in the nonvertebrate lineages.”

Stanhope et al. (2001) also refute the initial report, arguing that the results are not trustworthy because the method used is not the most appropriate since ‘‘phylogenetic reconstruction is critical to synthesizing, from the growing wealth of sequence data, a more comprehensive view of genome evolution.” Hence the authors conducted a phylogenetic assessment of 28 proteins (from the 113 corroborated set), discarding all of them as false positives ‘unlikely to be examples of direct HGT from bacteria to vertebrates.” [12]

This example clearly shows the controversy surrounding the possibility of lateral gene transfer to multicellular eukaryotes. A more recent publication, in 2015, sets out to resolve this issue. Using a phylogenetic approach with a much larger database of genomes and transcriptomes, Crisp et al. (2015) found 145 high-confidence foreign genes (not only of prokaryotic origin) that are present, and expressed, in the human genome, most of which are metabolic enzymes. This report, again led to a response article by Salzberg et al. (2017) that found ‘little or no evidence to support claims of horizontal gene transfer (HGT)”, clearly showing that this subject continues to spike heated debates.

Accordingly, there are much less reports regarding LGT in eukaryotes, most of them are either small-scale individual gene reports [13, 14, 15, 16, 17, 18, 19, 20, 21] or Endosymbiotic Gene Transfers (EGT), i.e. the intracellular gene transfer between organelles and the nucleus [22, 23, 9]. Whole genome LGT studies are even rarer (see [7] for a large scale prokaryotic study and the follow-up paper [24] challenging the magnitude of those findings), mostly because the bona fide method of LGT classification involves the evaluation of individual phylogenies for every gene, ideally containing only orthologous sequences, and such process is cumbersome and time-consuming to do manually, and error prone when done automatically on a large scale. Nevertheless, a few eukaryotic systematic LGT analyses have been published reporting significant numbers of LGT events, e.g in rumen ciliates [25], in *Entamoeba histolytica* [26], in *Trichomonas vaginalis* [27], in the diatom *Phaeodactylum tricornutum* [28], the fungal kingdom [29], the extremophile red algae *Galdieria sulphuraria* [30], and even in the multicellular *Hydra magnipapillata* [31] and in the tardigrade *Hypsibius dujardini* (where the first publication reports one sixth of the genes to be lateral transfers [32], and a subsequent analysis shows that those high values were in fact due to contaminations in the genome assembly [33]), showing the importance of such large-scale LGT studies for the understanding of the eukaryotic genome evolution.

*Piromyces sp*. strain E2 is a peculiar chytrid fungus that grows as a commensal organism in the gut of virtually all mammalian herbivores studied so far, e.g. cattle, goats, sheep and elephants to name a few [34]. It is an obligate anaerobic fungus belonging to the order Neocallimastigales, which comprises basal fungi that lack classic mitochondria. Instaed these organisms possess hydrogenosomes [35], i.e. Class 4 organelles of mitochondrial origin according to Müller’s classification [36]. These organelles have evolved from mitochondria and generate ATP via substrate phosphorylation, disposing of excess reducing equivalents via hydrogen-producing fermentation, and do not harbor a genome, cytochromes nor a membrane-associated electron transport chain.

Like all Chytridiomycota, Piromyces reproduces with flagellated motile spores (zoospores), but it displays several physiological peculiarities that set it apart from most other fungi. For example, Piromyces presents three enzymes of mitochondrial ancestry that have been retargeted, and are functionally active, in the cytoplasm – malate dehydrogenase, aconitase and acetohydroxyacid reductoisomerase [37, 38]. Also it exhibits a bacterial-type mixed-acid fermentation whereby, instead of producing ethanol from pyruvate via pyruvate decarboxylase and alcohol dehydrogenase (as in the alcoholic fermentation of yeast), it does so in the cytoplasm through the sequential action of pyruvate:formate lyase (PFL) + alcohol dehydrogenase E (ADHE) [35]. Moreover, Piromyces’ pyruvate catabolism in the hydrogenosome also uses PFL, just like the closely related fungus *Neocallimastix frontalis*, contrasting with other hydrogenosome-bearing organisms that usually use pyruvate:ferredoxin oxidoreductase (PFO) (for a comprehensive review see [38, 39]).

Herbivores feed mostly on cellulose, and are able to digest it by possessing massive fermentation vats inhabited by symbiotic cellulolytic microbes as part of their digestive tract [40]. Monogastric herbivores, like the elephant and the horse, rely on hindgut fermentation that takes place in their large caecum, while ruminants are foregut fermenters since the rumen is the first compartment of their digestive system. Despite this anatomical difference, the process of anaerobic fermentation that occurs in the hindgut is essentially identical to that which occurs in the forestomach of ruminants. Also, herbivores ferment the ingested food through the mutual action of symbiotic cellulolytic microorganisms. This gut microbiota is composed mainly by bacteria, archaea, unicellular eukaryotes (mainly ciliates) and anaerobic fungi, which are known to be key players for the hydrolysis of the plant cell-wall and its major component – cellulose [41].

Anaerobic fungi grow as saprophytes, and develop a mycelium that penetrates the fibrous plant material ingested by the herbivore [42], leading to a physical and enzymatic hydrolysis of plant cell-wall carbohydrates, delivering readily accessible nutrients to the host, and large amounts of hydrogen (H2) to the gut methanogenic community [43,44]. Since cellulose represents the most abundant reservoir of organic carbon in the biosphere, and anaerobic fungi secrete large amounts of cellulases and other enzymes into the environment, the gut fungal community represents a rich source of hydrolytic enzymes with great biotechnological potential, particularly related to direct fermentation schemes and biofuel production from lignocellulolytic biomass [44, 45]. In fact, several cellulase genes have been cloned from chytrid fungal strains such as *Neocallimastix patriciarum* [46], Orpinomyces PC-2 [47], and *Piromyces rhizinflatus* [48]. More recently, anaerobic gut fungi (and *Piromyces sp*. in particular) have also been the target of several studies focusing on using these fungi as microbial nutritional additive to enhance the digestibility of poor-quality lignocellulosic food, aiming to increase domestic cattle productivity [49, 50].

The highly symbiotic nature of Piromyces’ ecological niche – the herbivore gut – is known to favor the genetic exchange between microorganisms [51], and there have been reports of individual genes transferred from bacteria to diverse rumen eukaryotes [52, 53], and to the chytrid fungus *Orpinomyces joyonii* [13]. One particular large-scale LGT study has evaluated the transfer of genes from bacteria to rumen ciliates, finding an over-representation of genes involved in complex carbohydrate catabolism and anaerobic lifestyle [25]. Similarly, the genomes of the two parasitic species *Entamoeba histolytica* and *Trichomonas vaginalis*, show high numbers of candidate LGT (96 and 152 respectively) the majority of which encode metabolic enzymes related to carbohydrate and amino acid metabolism [27, 26]. Another study of the extremophile red algae *Galdieria sulphuraria*, shows a total transfer of 5% of its genes from prokaryotes [30], strongly supporting the thesis that LGT is an important evolutionary source of innovation, particularly significant in specific niches, like the gut, or environments with high temperatures, to which some bacteria and archaea have been adapted before the origin of some eukaryotic species, hence enabling the fast adaptation of eukaryotes to exploit new environments.

Given its evolutionary importance and biotechnological potential, the genome of *Piromyces sp*. strain E2 (isolated from an Indian elephant), has been sequenced as part of the Fungal Genomics Program of the DOE Joint Genome Institute (JGI), together with four other Neocallimastigomycota. Their recent genome publication focused on describing the set of proteins critical for fungal cellulosome assembly [54]. Remarkably, the authors report that 9-13% of the genes from the five gut fungi (no reports of specific values per organism) are more similar to bacterial than to eukaryotic genes. Individual results are only discussed for the domains of the proteins involved in fungal cellulosome assembly, namely the non-catalytic dockerin domains (NCDDs), hence leaving several open questions regarding the LGT events.

As such, we set out to elucidate which proteins were likely transferred to the *Piromyces sp*. E2 genome, from which bacterial donor species, and which functions were laterally acquired in Piromyces’ adaptation to the cellulose-rich and anaerobic environment of the elephant gut. Using sequence comparison and phylogenetic analyses, we have found 704 LGT candidates, representing nearly 5% of the *Piromyces sp*. orfeome (i.e. the complete set of open reading frames), mostly transferred from Firmicutes, Fibrobacteres, Bacteroidetes and Proteobacteria, closely following the microbial abundance reported for the herbivore gut. With respect to the functional analysis, the LGT candidate set includes proteins from 250 different orthologous groups, with a clear over-representation of genes belonging to the Carbohydrate Transport and Metabolism functional class. Moreover, we performed a graph density analysis on the metabolic pathways formed by the LGT proteins, showing that the connectivity between the LGT candidates would be very unlikely found if random enzymes were picked from the global KEGG metabolic map, meaning that the acquired proteins are non-random proteins that fit within Piromyces metabolic network.

## Results

In this study we performed a detailed large-scale lateral gene transfer (LGT) analysis of the genome from the chytrid fungus *Piromyces sp*. strain E2. For this we evaluated its 14648 predicted proteins, and mapped the LGTs occurring between prokaryotes and this anaerobic eukaryote. Briefly, we started by ranking the proteins according to their LGT likelihood using the distribution of the BLASTP hits as a proxy for phylogenetically atypical proteins (see the Methods section for a detailed explanation). Subsequently, possible transfers from other eukaryotic species were filtered out since we chose to focus our analysis on investigating the transfer of bacterial and archaeal genes to the eukaryotic genome of Piromyces. Finally, we performed an individual phylogenetic evaluation for each of the preliminary candidates, ending up with a set of 704 likely LGT candidates, representing 4.8% of its proteome.

### Functional analysis of the LGT candidates

#### Over-representation of carbohydrate transport and metabolism

To determine the functions of the proteins laterally transferred we have firstly assigned the 704 LGT candidates to their respective COG functional categories. Their distribution is clearly enriched for *Metabolic* function classes (shown in green in Figure 1), with the *Carbohydrate transport and metabolism (G)* category alone representing more than one quarter of the total (27.41%). *Cellular processes and signaling* (in orange) represent about 13%, and *Information storage and processing* (in purple) a residual 4.4% of the total. Despite the large fraction of *Poorly characterized functions* (in blue) there is still a part of *General function prediction* (the wavy wedge in category R) whose annotations contain the keyword “cellulosome”.

**Figure 1.**
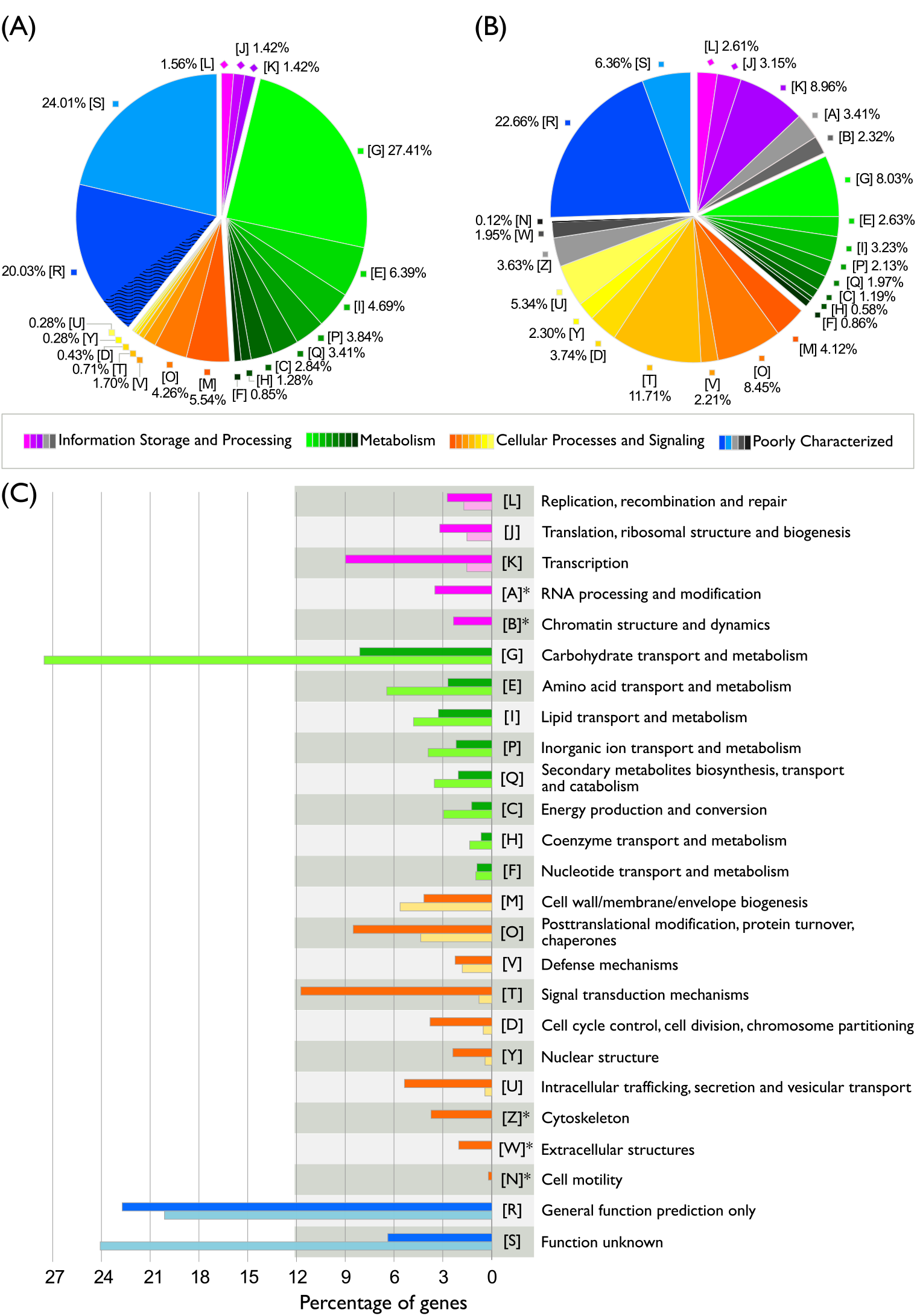
Function category distribution in the LGT candidate list (A), in the whole genome (B), and the comparison between both (C). In 5A the wavy stripes in category [R] indicate the percentage of proteins whose general function prediction includes the word cellulosome. In 5B the gray fractions of the pie chart represent the functional classes that are not represented in the LGT candidate set. (Note that the percentages in 5A and 5B do not add up to 100% since some proteins belong to more than one functional category, namely 1180 proteins out of 8626 in 5A and 70 out of 704 proteins in 5B). In 5C the darker bars represent the functional classes distribution in the whole genome and the lighter bars the distribution of the LGT candidates’ functions. Asterisks indicate the 5 categories that are not represented in the LGT candidate set.

The cellulosome is a large complex of cellulolytic enzymes, consisting of a central organizing protein, named scaffoldin, which binds the enzymes by its cohesin sites, and of individual catalytic components: Carboxymethylcellulases (endoglucanase), Cellobiohydrolases (exoglucanase) and B-glucosidases, which in turn require a dockerin domain for incorporation into the complex [55, 56]. Together they degrade and solubilize cellulose, hemicellulose and other plant cell-wall polysaccharides. The presence of this keyword (cellulosome) plus the other cellulosome-associated enzymes in the functional annotations of 71 top-ranking candidate LGTs, shows the importance of LGT for the carbohydrate transport and metabolism in Piromyces (Table 1), confirming the previous report that some catalytic domains from fungal cellulosomes originated via lateral gene transfer from gut bacteria [54]. In contrast, the most prevalent function found in Piromyces’ genomic background (Figure 1 C) is *Cellular processes and signaling* (43.57%). In fact, the combined 8 metabolic classes in the whole genome represent only circa 20% of the total (Figure 1 B), making the nearly 50% metabolic classes found in the LGT candidate set stand out.

**Table 1.** Top-20 functions found in the LGT set (excluding the category S — function unknown). The most frequent functional description among all the proteins belonging to the same Orthologous Group is displayed.

| # | Functional Class | Function Description |
| --- | --- | --- |
| 38 | Poorly Characterized [R] | Predicted hydrolases or acyltransferases ($\alpha/\beta$ hydrolase superfamily) |
| 34 | Metabolism [G] | ABC-type sugar transport system, periplasmic component |
| 34 | Metabolism [G] | Endo-1,4-β-xylanase (EC 3.2.1.8) |
| 24 | Poorly Characterized [R] | Cellulosome enzyme, dockerin type I precursor |
| 21 | Metabolism [M] | Spore coat assembly protein |
| 15 | [GEPR] | Permeases of the major facilitator superfamily |
| 13 | Metabolism [G] | Endoglucanase |
| 11 | Poorly Characterized [R] | Predicted phosphohydrolases |
| 10 | Metabolism [G] | Acetylxylan esterase-like protein |
| 10 | Metabolism [I] | Esterase/lipase |
| 9 | Metabolism [Q] | Poly(3-hydroxybutyrate) depolymerase |
| 9 | Poorly Characterized [R] | Aldo/keto reductases, related to diketogulonate reductase |
| 6 | Metabolism [C] | Uncharacterized oxidoreductases, Fe/dependent alcohol dehydrogenase family |
| 6 | Metabolism [G] | Predicted xylanase/chitin deacetylase |
| 6 | Poorly Characterized [R] | Predicted peptidase |
| 5 | Metabolism [G] | β-mannanase |
| 5 | Metabolism [I] | Carboxylesterase type B |
| 5 | [MG] | Predicted nucleoside-diphosphate-sugar epimerases |
| 5 | Metabolism [P] | Enterochelin esterase and related enzymes |
| 5 | Metabolism [Q] | O-Methyltransferase involved in polyketide biosynthesis |

These results are in agreement with the hypothesis that horizontal transfer occurs frequently among symbiotic communities; but the acquired genes will be retained only when they confer a fitness increase to the receiving species [2, 57]. Given the environmental conditions found in the gut of herbivores where energy-rich complex carbohydrates are an abundant resource, one could expect that gaining genes related to carbohydrate transport and metabolism would increase the metabolic toolkit available to the host species, hence conferring it a major competitive advantage. In fact, a similar trend has been previously reported for the ciliate rumen community, where 75% of predicted horizontal gene transfers were related to metabolism and 39% alone devoted to carbohydrate transport and metabolism [25].

The importance of LGT of genes related to carbohydrate metabolism has also been observed in the anaerobic parasitic species *Entamoeba histolytica* and *Trichomonas vaginalis*, where it appears to have exerted a major impact in the range of substrates available for energy conversion, enabling the usage of readily available sugars other than glucose, like fructose and galactose [26, 27]. Our LGT results strengthen the evidence that indeed the eukaryotic adaptation to the cellulose-rich niche of the herbivore gut has been fostered by the lateral acquisition of genes from cellulolytic bacteria.

#### Glycosyl-hydrolases and ABC-type sugar transport enrichment

After evaluating the functional classes that grouped most LGT candidate proteins, we studied the predicted functional annotation of individual proteins to discover which particular functions were over-represented in the LGT candidate set.

We have found an enrichment of sugar transporters (which is quite an interesting finding since the eukaryotic membrane is considerably different from the prokaryotic one) and fibrolytic cellulosome enzymes, as well as several glycosyl hydrolases: endoxylanases, endoglucanases, acetylxylan-esterases and beta-mannanases) (Table 1).

Moreover, 30% of these enzymes (212 in total) are predicted, by TargetP [58] to be secreted (data not shown), consistent with the mechanism of extracellular enzymatic hydrolysis of cellulose undertaken by gut anaerobic fungi. Unlike bacteria that possess cell-wall attached cellulosomes, cellulolytic chytrids secrete enzymes into the environment where they compose a multiprotein cellulose-binding complex that has been shown to convert crystalline cellulose exclusively into glucose [59].

#### Hydrogenosomal proteins present in the LGT candidate set

Since *Piromyces sp*. E2 is one of the few species known to harbor a hydrogenosome, we investigated the occurrence of putative hydrogenosomal proteins in our LGT candidate set. The putative hydrogenosomal proteome list was constructed by combining the results from the following procedures: (i) all predicted ORFs were run through TargetP [58] and WoLF PSORT [60] to find organellar predictors; (ii) all proteins were searched against six biochemically determined mitochondrial proteomes (*Arabidopsis thaliana, Chlamydomonas reinhardtii, Homo sapiens, Mus musculus, Tetrahymena thermophila* and *Saccharomyces cerevisiae*); (iii) then all orfs were searched against a list of enzymes known to be involved in anaerobic biochemistry so that we would not miss unexpected anaerobic processes; (iv) finally the sequences were searched against predicted/confirmed mitosomal and hydrogenosomal proteomes from *Giardia lamblia, Entamoeba histolytica* and *Trichomonas vaginalis*. (The list resulting from the union of these criteria is available online as Supplementary data 1 at: http://doi.org/10.6084/m9.figshare.7173020.v1).

We found an overlap of 26 LGT candidates with the putative hydrogenosomal proteome set (Table 2). Among these, several pivotal functions in hydrogenosomal metabolism are represented, namely the Pyruvate-formate lyase (PFL) family (further details in the Discussion), Glutathione peroxidase, Alcohol dehydrogenases and ABC transporters. This fact suggests that some LGT proteins that might be targeted and functioning in the hydrogenosome have been acquired after the establishment of the proto-mitochondrion, since none of these proteins are traced back to the alpha-proteobacteria (the group that gave rise to the mitochondrion and related organelles).

**Table 2.** Lateral Gene Transfer candidates that overlap with the set of putative hydrogenosomal proteins.

| # | Functional Class | Function Description |
| --- | --- | --- |
| 5 | Cellular Processes and Signaling [O] | Glutathione peroxidase (4);<br>Membrane protease subunits, stomatin/prohibitin homologs (1). |
| 3 | Metabolism [C] | Pyruvate-formate lyase (2);<br>Uncharacterized oxidoreductases, Fe-dependent alcohol dehydrogenase family (1). |
| 2 | Metabolism; Poorly Characterized [IQR] | Dehydrogenases with different specificities (related to short-chain alcohol dehydrogenases). |
| 3 | Metabolism [I] | Esterase/lipase (2); Long-chain acyl-CoA synthetases (AMP-forming) (1). |
| 2 | Metabolism [E] | Tryptophanase (1); Glutamate decarboxylase and related PLP-dependent proteins (1). |
| 2 | Cellular Processes and Signaling [V] | ABC-type multidrug transport system, ATPase and permease components (1). |
| 1 | Metabolism [G] | 6-phosphofructokinase. |
| 1 | Metabolism [H] | Aspartate oxidase. |
| 1 | Metabolism [F] | Uracil phosphoribosyltransferase. |
| 1 | Cellular Processes and Signaling [M] | dTDP-4-dehydrorhamnose reductase. |
| 3 | Poorly Characterized [R] | Predicted hydrolases or acyltransferases ($\alpha/\beta$ hydrolase superfamily) (1);<br>Zn-dependent hydrolases, including glyoxylases (1);<br>Predicted amidohydrolase (1). |
| 2 | Poorly Characterized [S] | No-annotation. |

### Metabolic pathway analysis of the LGT candidates

To visualize function interactions between the LGT candidate proteins we used iPATH2 [61] to highlight all Piromyces’ proteins in the cellular metabolism pathway map (Figure 2). Blue edges represent vertically inherited metabolic enzymes present in the genome, and red edges mark the candidate lateral gene transfer orthologous groups (OG). The width of the edge is proportional to the number of LGT proteins belonging to that OG. From the 250 total OGs represented in our LGT dataset, 85 were successfully mapped to the metabolic pathway map. A relevant portion of the laterally transferred enzymes seems to be functionally related, since continuous portions of metabolically connected pathways appear to have been laterally acquired.

**Figure 2.**
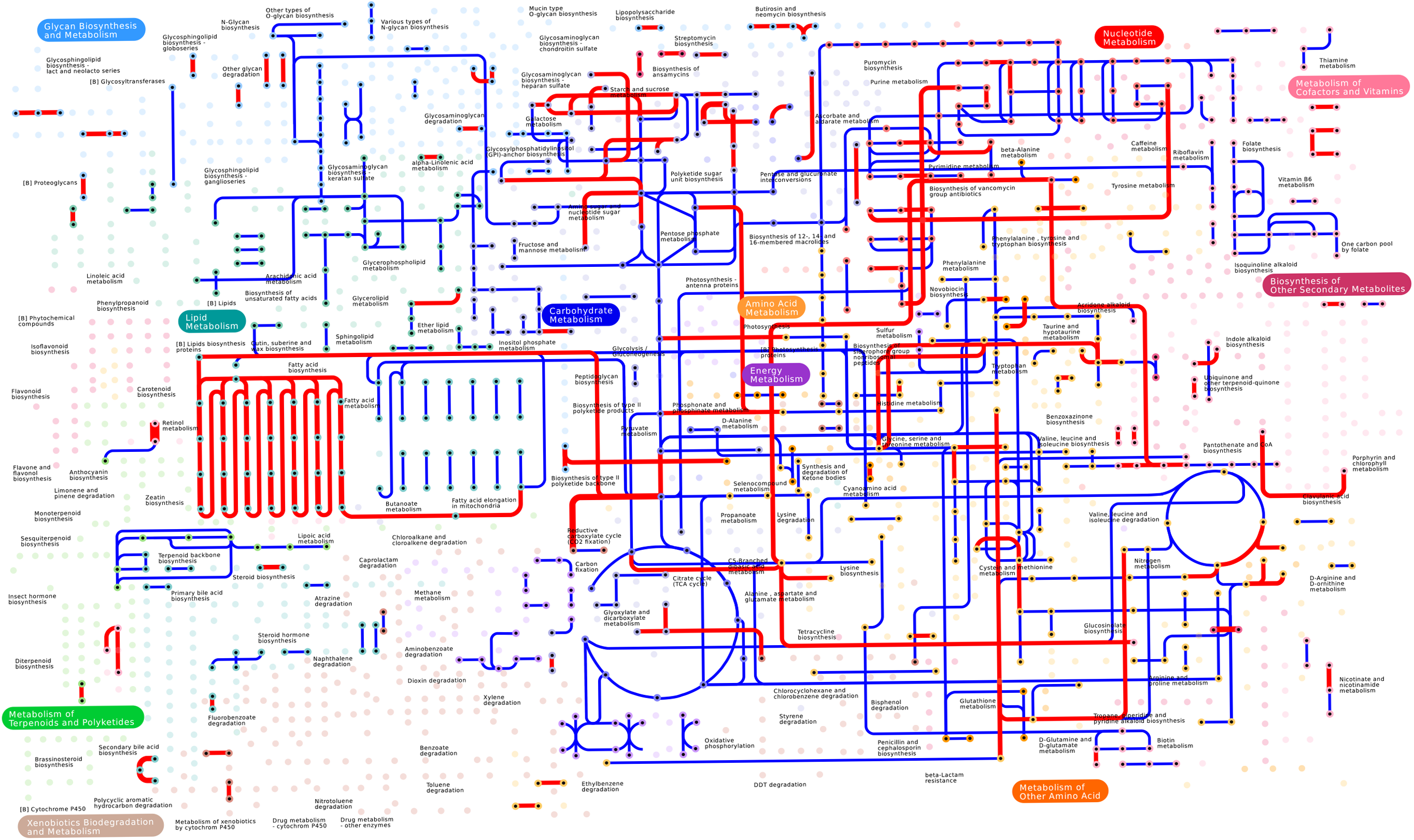
Cellular metabolism pathway map. iPATH2 blue edges represent the proteins present in the *Piromyces sp*. E2 genome and red edges highlight the lateral gene transfer (LGT) candidate proteins for which there are Orthologous Group (OG) or KEGG annotations. The edge width is proportional to the number of LGT instances mapping to the same edge. The significance of the LGT candidates connectivity (red edges) has been studied and shown to be significant when compared to the overall KEGG map (grey edges), and not significant when compared to the whole Piromyces metabolism (blue edges). (More details on the section for pathway cohesion analysis.)

#### Pathway cohesiveness

In order to find if the visually apparent connectivity of the LGT metabolic genes is significantly different from what could be expected from randomly choosing enzymes, either from the Pan-metabolic map (the general map of all known metabolic reactions) or from the Piromyces background metabolism (which is a subset of the Pan-metabolism), we calculated the densities of the LGT and Piromyces’ metabolic graphs, and compared them to the density distribution of 1 million random graphs generated by sampling edges from the appropriate background graph.

As expected, all the graphs display very low density values (Table 3), since metabolic pathways tend to be linear (1 edge between each 2 nodes) and not very strongly connected networks. Accordingly, the least dense graph is the whole metabolism (1.43*E* – 3), followed by the Piromyces’ metabolism (5.32*E* – 3) and the highest density was found in the LGT graph (7.34*E* – 3). Altogether, these results corroborate the fact that the whole metabolic map (displaying all known individual pathways) will inevitably be more loosely connected, as a whole, than a subgraph representing the metabolic pathways of a particular organism. Moreover, these values accurately grasp the visual perception of connectivity associated with each map (Figure 2).

**Table 3.** Graph density values for observed metabolic graphs and average density values estimated from the distribution of 1 million random samples.

|  | Density | Metabolic Graph Description |
| --- | --- | --- |
| Observed Values | $1.43E - 3$ | KEGG Pan-metabolic map (1939 edges) |
| | $5.32E - 3$ | *Piromyces* metabolism (868 edges) |
| | $7.34E - 3$ | Lateral Gene Transfer (LGT) metabolism (136 edges) |
| Estimated from Distribution | $4.34E - 3$ | LGT-like random graph sampled from Pan-metabolic map (136 edges sampled from 1939 edges) |
| | $7.55E - 3$ | LGT-like random graph sampled from *Piromyces* metabolism (136 edges sampled from 868 edges) |

When comparing the observed density values with the random density distributions (Supplementary figure 1 available online at: https://doi.org/10.6084/m9.figshare.7173020.v1)), two main conclusions stand out: (i) The Piromyces’ LGT metabolism has a graph density of 7.34*E* – 3, which is significantly higher than the average density of 4.34*E* – 3 found by randomly choosing 136 edges from the Pan-metabolism graph (which has 1939 edges). In fact, in 1 million random graph simulations we found no single value higher or equal to the LGT’s observed density. In other words, the observed LGT density has a p-value < 1/ 1E6, meaning that this cohesion between the LGT candidates would be unlikely found if it were randomly picked from the Pan-metabolism. This observation shows that the LGT genes form cohesive non-random metabolic modules within the scope of the Pan-metabolic map. (ii) When randomly chosen from the Piromyces’ background metabolism, the average density of 7.55*E* – 3 is not significantly different from the calculated LGT graph density (7.34*E* – 3), meaning that the laterally transferred proteins do not form particularly cohesive metabolic pathways within the Piromyces’ metabolism. Accordingly, it is more likely that the LGTs that became fixed in the population have been plugged into Piromyces’ core metabolism, as opposed to being transferred in bulk as metabolic modules.

### 3. The major donor taxa mirror the bacterial abundance in the herbivore caecum

To answer the question: “Where do the laterally transferred proteins come from?”, we evaluated the maximum likelihood phylogenies calculated for the 240 high-confidence candidate LGT set. This selection assures that we include in the donor taxa analysis only reasonably well-supported branches. Eleven different bacterial phyla were found, while only the Euryarchaeota phylum was represented from the Archaea domain (Figure 3).

**Figure 3.**
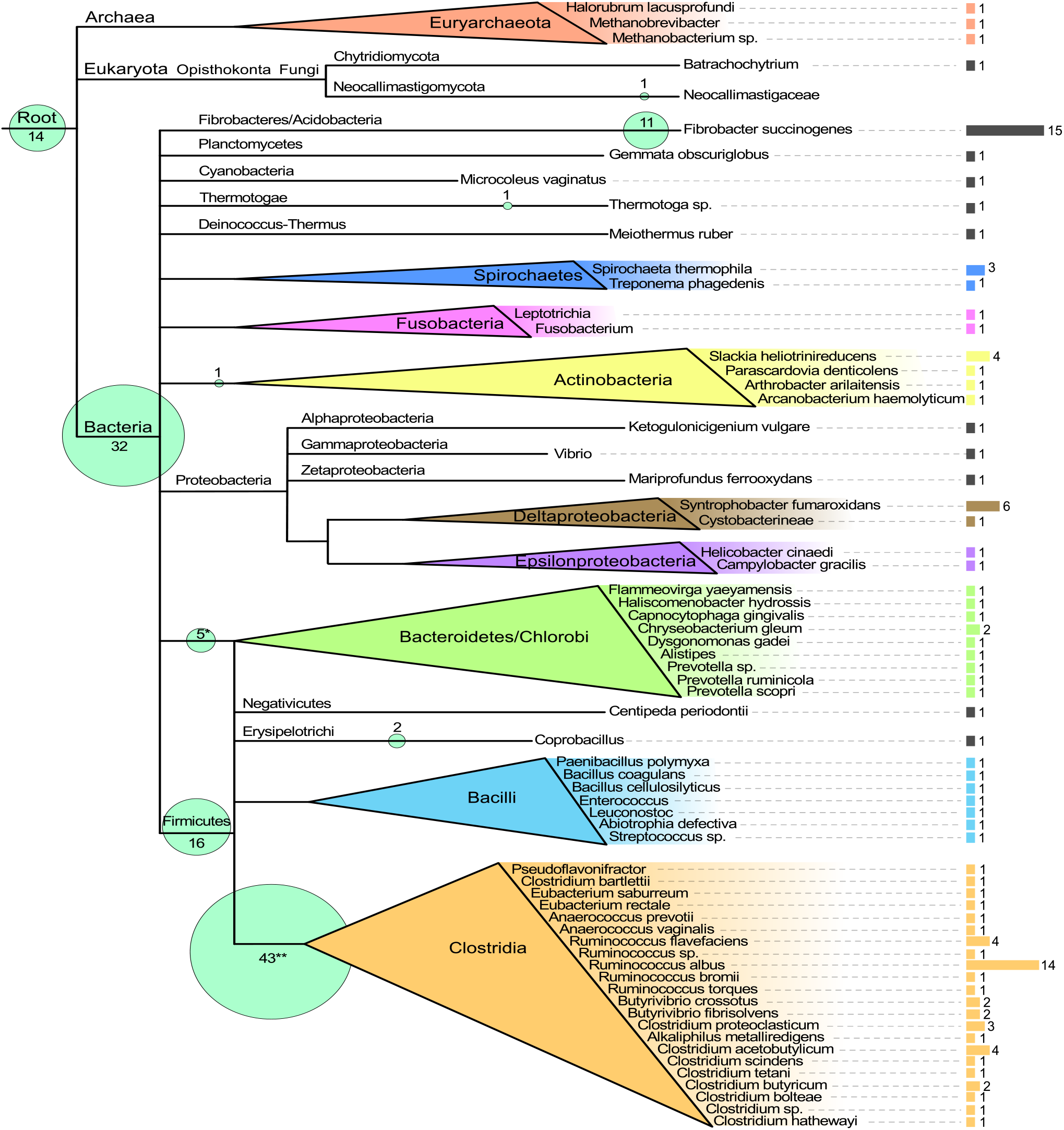
Trimmed ‘‘Tree Of Life” showing the donor taxa that contributed to the 240 high-confidence lateral gene transfer (LGT) candidate list. Genus or species-level taxa are displayed in the leaves and the number of times that this taxon appears as last-common-ancestor in the high-confidence LGT candidate set is shown by the size of the lateral bars. Other non-species last-common-ancestor taxonomic rankings are marked by inner-branch bubbles, with the respective values indicated next to it. (Asterisks indicate bubbles that contain more than one closely related taxa, as follows: (*) 2 Bacteroidetes + 1 Bacteroidales + 1 Prevotella + 1 Flavobacteriaceae; (**) 5 Clostridia + 29 Clostridiales + 2 Clostridium + 2 Blautia + 2 Butyrivibrio + 3 Ruminococcus).

The most recent characterization of the gastrointestinal bacterial microbiota of a monogastric non-ruminant herbivore (the horse’s gut) reported that there are marked differences between individual gut compartments, with the caecum (the compartment where the fermentation takes place) showing the Firmicutes as the most abundant phylum (ca 58%, with Clostridia being the most frequent Class), followed by the Verrucomicrobia (ca 10%), the Proteobacteria (ca 8%), Fibrobacteres (ca 6%), Spirochaetes (ca 5%) and Bacteroidetes (ca 5%) [62]. Appropriately, the taxonomic distribution of our high-confidence LGTs closely mirrors the microbial abundance found in the horse’s caecum, with the most represented phylum being the Firmicutes (48.3% of the transfers (116 out of 240, of which 89 are from the Clostridia class), followed by the Fibrobacteres (10.8%), Bacteroidetes (6.3%) and Proteobacteria (5.0%) (Figure 3).

Regarding the difference in the relative raking of Fibrobacteres (second in our LGT donor set compared to fourth in the horse’s gut abundance), one can speculate that it might be either due to (i) particular differences in the relative-abundance of each Phylum in the elephant; or (ii) because the Fibrobacteres’ genes confer a greater metabolic advantage to Piromyces, hence being more often successfully retained after lateral transfer.

The individual bacterial species that contributed the most to our LGT set are the two best described primary degraders of plant fiber: *Fibrobacter succinogenes* – involved in 15 LGTs and *Ruminococcus albus* – a Clostridium involved in 14 LGTs. These two anaerobic species, given their efficient cellulolytic activity, have been extensively studied for biotechnological applications, and are among the most important cultured cellulose degrading bacteria found in the herbivore gut [63, 64]. Interestingly, they present two alternative strategies of hydrolyzing cellulose. *Ruminococcus albus* is a gram-positive bacterium belonging to the Clostridia group, well known for its cell-wall attached cellulosome, which enables the bacterium to adhere to the substrate and enzymatically degrade it [65]. In contrast, *Fibrobacter succinogenes*, which appears to use cellulose as its sole energy source, is a gram-negative bacterium that does not use a cellulosome nor does it produce high extracellular titers of cellulase enzymes like other cellulolytic microorganisms [66]. Despite the many proposed theories, its exact hydrolysis mechanism remains unknown and a topic of active research [66].

An analysis of the identity of the LGT candidates originating from these two species, revealed that out of the total 29 proteins transferred, 21 belong to the functional category *Metabolism*, 5 are only annotated with the general function *cellulosome*, and only 3 belong to other function classes (Table 4). Nearly all proteins with specific metabolic functions are annotated to be enzymes related to polysaccharide hydrolysis, e.g. beta-xylanases, beta-mannanases, endoglucanases, and cellulose binding, showing that indeed the metabolic genes, particularly the ones that expand the capability of using cellulose and its derivatives as a carbon source in such a niche, are the ones that get fixed after their transfer from cellulose degrading bacteria.

**Table 4.**
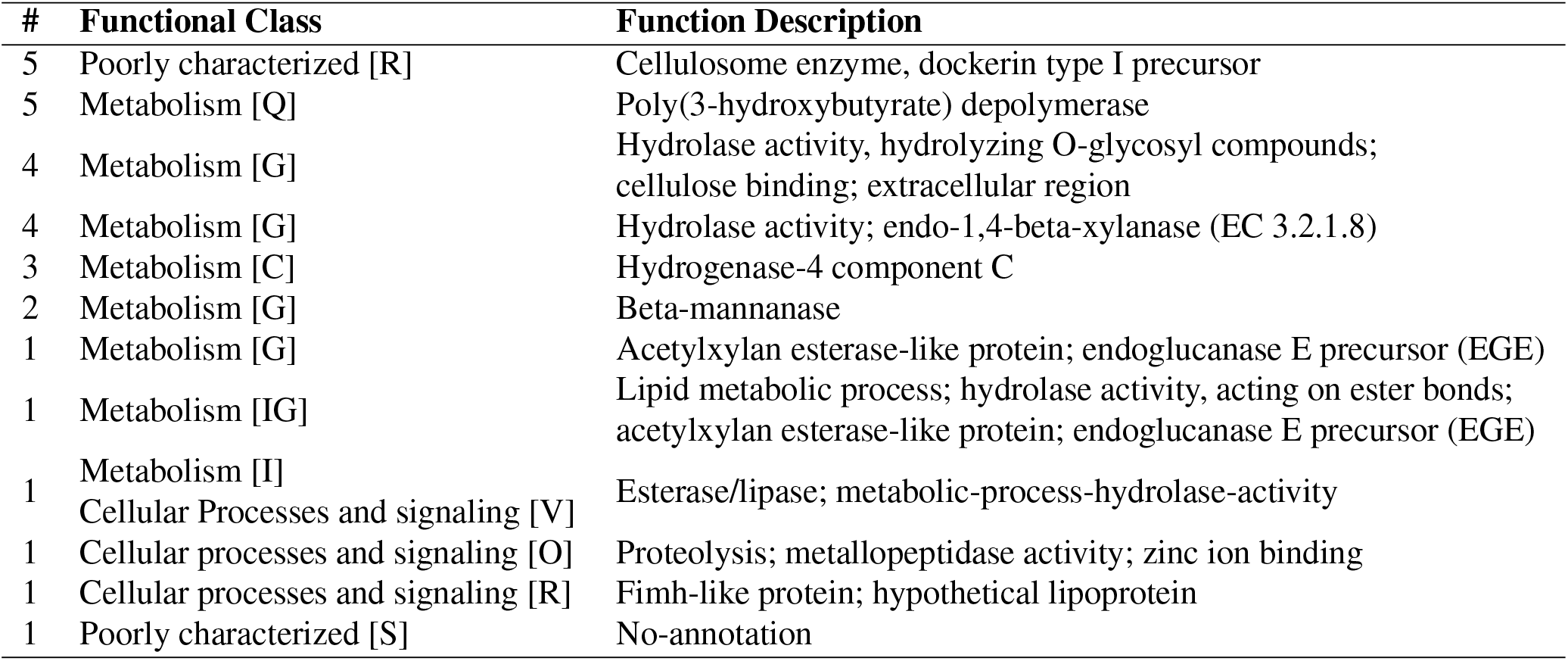
Functional categories and functional description of the 29 Lateral Gene Transfer candidates from *Fibrobacter succinogenes* and *Ruminococcus albus*.

### 4. Under-representation of archaeal LGT candidates

Remarkably, despite the fact that approximately 3% of gut microbes are autotrophic methanogenic archaea, only two LGT candidate proteins seem to have an archaeal origin: one oxidoreductase (functional class H) from Methanobrevibacter, and one methyltransferase (class R) from Methanobacterium. (There is a third candidate, classified as a transporter from category G, but this is highly likely to be an artifact due to the poor coverage of Archaeal genomes sequenced, since *Halorubrum lacusprofundi* is a halophilic bacterium living in salty water and not in the herbivore gut.) This is in line with the recurrent observation that horizontal transfers between eukaryotes and archaea are less common [67, 25], with the notable exception of *Galdieria sulphuraria*, whose whole genome sequencing revealed a significant number of Archaea to Eukaryota gene transfers [30]. However, this observation cannot be dissociated from the fact that Galdieria is an extremophile algae that inhabits a hot, toxic and acidic environment populated mostly by Archaea. Accordingly, it would be expected that any transfers to Galdieria would originate from Archaeal prokaryotes sharing the same ecological niche.

Three possible factors might explain the observed scarcity of LGT between Archaea and Eukaryota. Firstly, most established LGTs belong to operational gene classes, probably because these are the ones that allow a faster adaptation of the organism, leading to an effective fitness increase that fixates the transferred gene in the population. Also, the modular nature of operational genes (i.e. the fact that they are part of smaller and less complex systems than informational genes) makes them more amenable to frequent horizontal gene transfer – the so called complexity hypothesis [68]. Finally, the well-documented bacterial origin of the eukaryotic operational genes [69], challenges the individual plugging-in of archaeal enzymes into the eukaryotic biochemical framework. This is mostly because the archaeal metabolic enzymes, and to some extent their pathways too, are substantially different from the ones employed by bacteria and eukaryotes [70], making the introduction of singular novel enzymes irrelevant to the eukaryotic metabolic toolkit, and hence usually not observed.

## Discussion

We set out to evaluate the impact of a symbiotic environment on the genetic evolution and metabolic diversification of the commensal gut fungus *Piromyces sp*. strain E2. Through our analyses, we have revealed that 5% of its genome has been transferred from Bacteria, which interestingly fits with the values published for other free-living eukaryotes that live in “extreme” niches, namely the 5% found in the extremophilic red algae *Galdieria sulphuraria* [30] and the 7.5% reported for the diatom *Phaeodactylum tricornutum* [28].

Accordingly, lateral gene transfer seems to have been an important force driving the environmental adaptation of the Piromyces’ genome to its highly specialized habitat. In fact, nearly 50% of the acquired genes code for enzymes involved in metabolizing the abundant sugars present in the herbivore gut, namely ABC-type sugar transporters, xylanases, endoglucanases and cellulosome-related subunits. Most of these proteins have been acquired from Clostridia, a class of anaerobic Firmicutes that are well-known gut inhabitants. It is evident from our results that the transfer of foreign genes closely follows the diversity of the gene pool, clearly revealed by the large overlap between the over-represented taxa in our LGT dataset and the relative abundance of those taxa in the gut microbiota. This fact, together with the non-random metabolic connectivity displayed by the LGT candidate set when compared to the whole metabolic map, clearly shows that the enzymes acquired via lateral transfer have been effectively plugged into Piromyces’ background metabolic pathways, potentially granting significant adaptive strength to this commensal fungus.

Piromyces’ adaptation to living anaerobically seems also to have been influenced by LGT. The overlap between the putative hydrogenosomal proteome and our high-confidence candidates is small (26 proteins) but it reveals 11 enzymes connected to metabolic functions (Table 2) which could also be functioning inside hydrogenosomes. One particularly interesting candidate is the pyruvate:formate lyase (PFL) (or formate C-acetyltransferase), which is known to be the major player in the pyruvate catabolism (decarboxylation of pyruvate to acetyl-CoA) taking place inside the hydrogenosomes from chytrid fungi [35]. This circumstance is rather unusual since PFL is a characteristic enzyme of facultative anaerobic Enterobacteria and Firmicutes that perform mixed-acid fermentation (ethanol is generated from acetyl-CoA and not from pyruvate), whereas eukaryotes (in particular trichomonads) mostly use pyruvate:ferredoxin oxidoreductase (PFO) instead [35]. However Boxma and colleagues have experimentally demonstrated that *Piromyces sp* E2 indeed exhibits a bacterial-type mixed-acid fermentation using PFL for the degradation of carbohydrates, and that it possesses an alcohol dehydrogenase E (ADHE), which depends on acetyl-CoA for the production of ethanol [35]. Additionally, the authors performed a phylogenetic analysis of ADHE where they show that “the evolution of ADHE appears to have involved at least one horizontal transfer event to the eukaryotes from the Firmicutes”. However, the lack of a PFL sequence precluded the analysis of its origin: “ (…) PFL might either be a eukaryotic relic or also be acquired by lateral gene transfer” [35]. Here, we answer this question, hence further enlightening the origin of chytridial central anaerobic metabolism by providing evidence that its PFL has been laterally acquired from Firmicutes, most likely from the Clostridia class (Figure 4).

**Figure 4.**
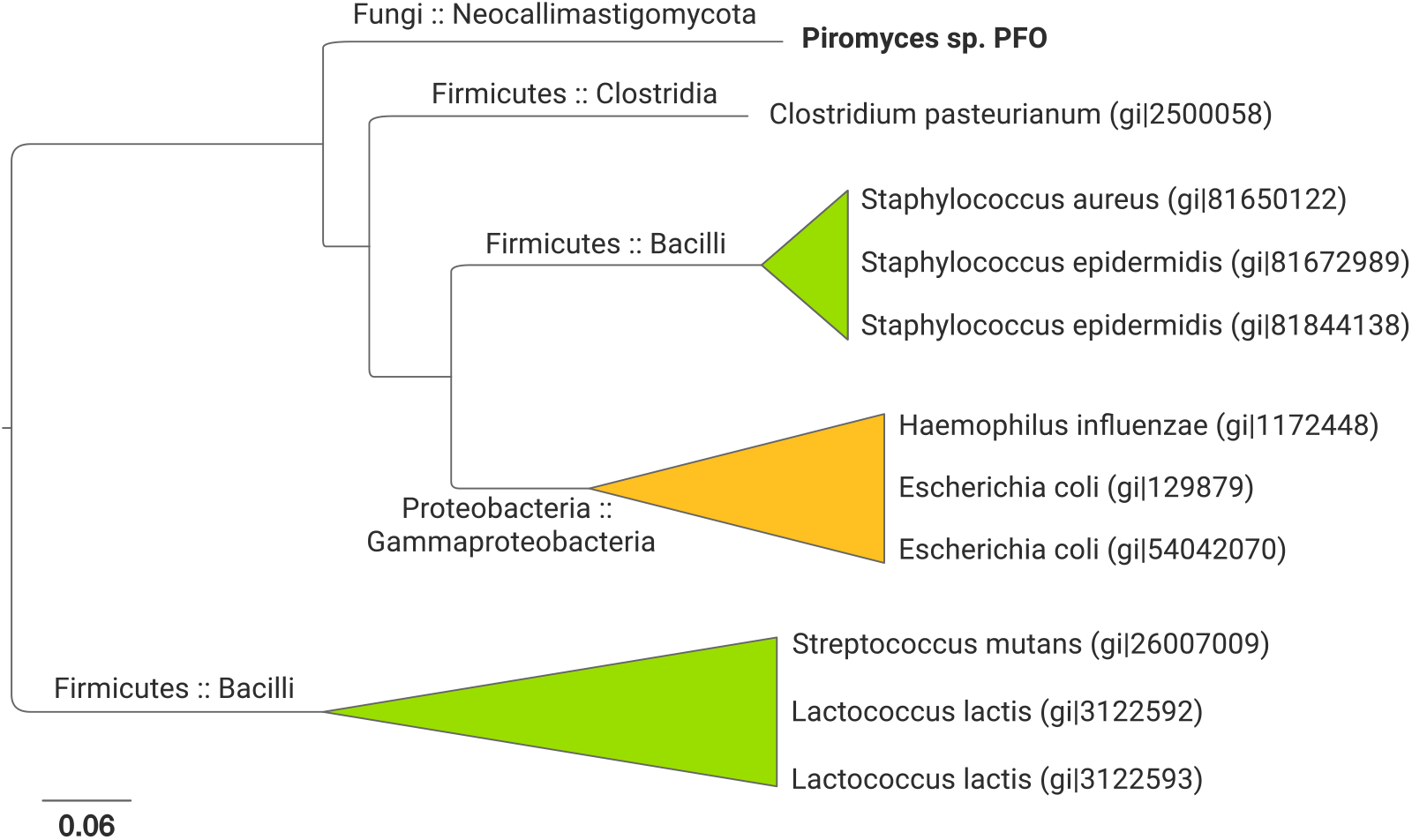
*Piromyces sp*. E2 pyruvate:formate lyase phylogeny. This unrooted NJ tree shows that only bacterial sequences are retrieved from the SwissProt database (E-value lower than 0.001) using as query the putative PFL sequence from Piromyces (GenBank:OUM56758.1). Moreover, Firmicutes are the closest sequences, with the Clostridia class displaying the smallest distance to our candidate sequence. A similar BLASTp search using NCBI’s refseq_protein database (Release 84) retrieves 1010 total sequences, all from Bacteria, with the closest 364 sequences (as indicated by the E-value) being from the Firmicutes phylum (data not show).

Overall, *Piromyces sp*. E2 represents not only an important source of biotechnological potential, but also an important evolutionary link, pivotal for our understanding of the mechanisms of eukaryotic genetic evolution and adaptation to confined symbiotic niches. Piromyces’ adaptation to living anaerobically and in the a cellulose-rich environment has been undoubtedly fostered by the acquisition of foreign genes from bacterial neighbors, showing the global importance of such evolutionary mechanisms for successful eukaryotic thriving in such competitive harsh environments.

## Methods

### Data retrieval

We have retrieved and analyzed the “Gene Catalog proteins” dataset from the *Piromyces sp*. strain E2 genome website (http://genome.jgi.doe.gov/PirE2_1/PirE2_1.home.html). It contains 14648 protein sequences derived from the automatic translation of the gene sequences contained in the ‘‘Filtered Models” dataset, meaning that these sequences were derived from the models representing the best gene model for each locus. The maximum sequence size is 6623 amino acids and the minimum is 49, with an average length of 386 residues.

### Lateral Gene Transfer candidate ranking

In order to automatically identify good lateral gene transfer (LGT) candidates, we started by using the DarkHorse (v1.4) software that applies a statistical method for discovering and ranking phylogenetically atypical proteins on a genome-wide scale, using relative levels of sequence similarity (for a detailed explanation of the algorithm, see [71, 72]).

Briefly, first it selects potential orthologs for all proteins of the genome of interest using BLASTP [73] hits for the proteome set against NCBI’s nr database. Then, based on the taxonomy of these matches (using NCBI’s Taxonomy database), it calculates a lineage probability index (LPI) score for each individual protein, which is then used to rank potential LGT candidates. Low LPI scores indicate lower phylogenetic relatedness between the query protein and its closest non-self BLASTP hits, hence making good LGT candidates.

First, the 14648 proteins in the dataset were used as query for a local BLASTP (v2.2.25) [73] search against the nr database (500 hits and e-value 0.05). These results were then used as input for a DarkHorse search, which was run with default parameters plus the following two settings: list as “self-hits” all taxa bellow the rank Order, i.e. the phylogenetic granularity of the results was set so that “older” LGT events would be identified by excluding from the search the following terms: Chytridiales, Chytridiomycetes, Chytridiomycota, Neocallimastigaceae, Neocallimastigales, Neocallimastigomycetes, Neocallimastigomycota and Piromyces; and 0.1 filter threshold, meaning that after removing “self-hits”, only results with BLASTP bitscore within 10% of the bitscore from the best-hit will be included in the subsequent LPI calculations, i.e. only well conserved hits were used. DarkHorse found 12503 proteins with non-self hits, for which it calculated LPI scores (Figure 5).

**Figure 5.**
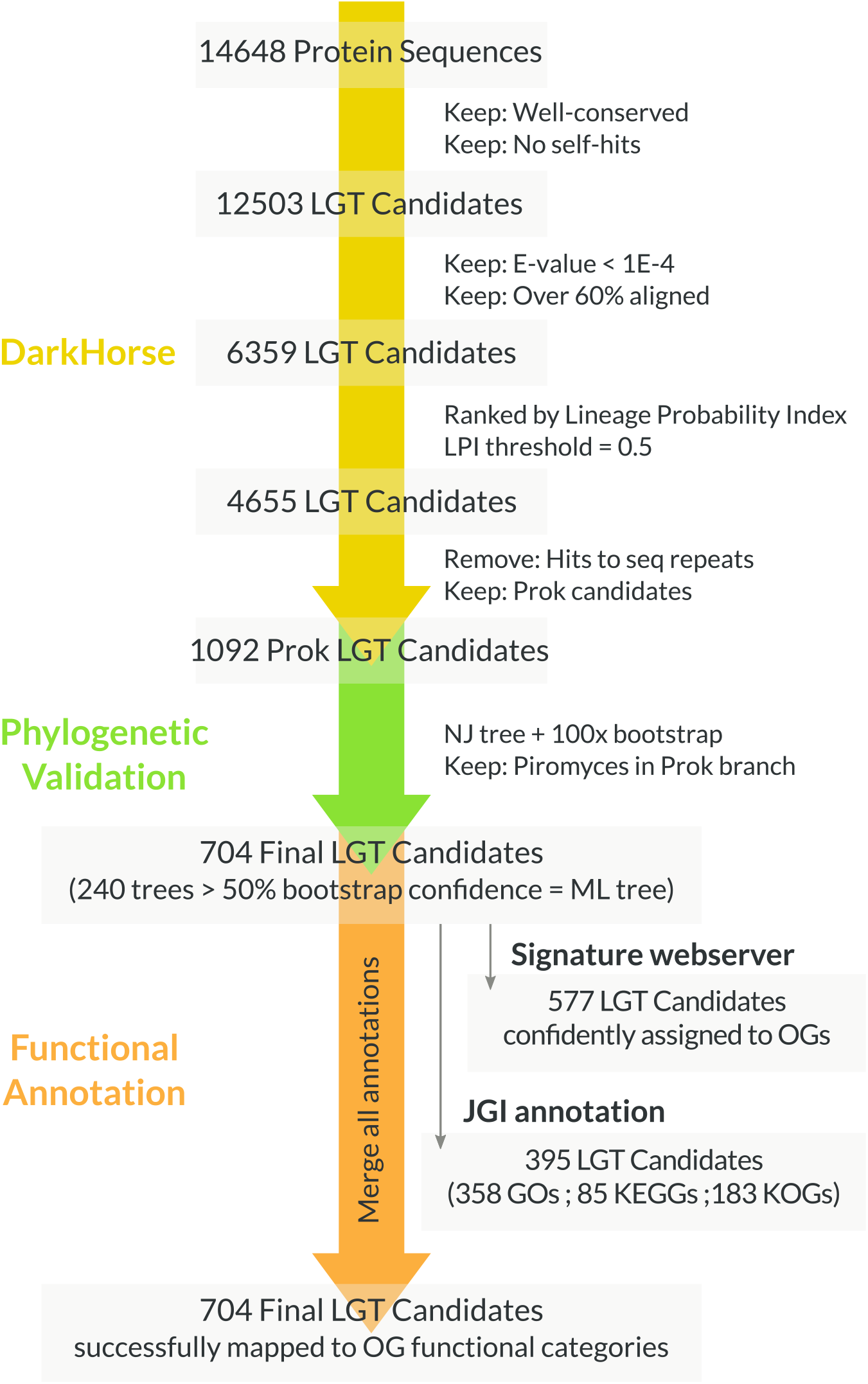
Data analysis pipeline. General schema summarizing the three main stages of data processing and analysis, namely the lateral gene transfer candidate list generation using the DarkHorse algorithm, the phylogenetic validation, and the final functional annotation.

After careful inspection of the results, it became clear that before ranking the LPI scores, the data required some filtering to remove misfit data. Accordingly two initial filters were applied: (1) remove hits with E-value above 1*E* – 4 to eliminate ill-aligned sequences; and (2) keep only the results which are over 60% aligned to the query, intending to filter out partial hits restricted to particular domains, or limited portions of proteins, that do not contain information for phylogenetic analyses. These filters effectively halved the number of initial results from 12503 to 6359.

Next, the 6359 candidates were ranked by LPI score. Following DarkHorse guidelines, to adjust the LPI threshold value to be used as cut-off for the ranking we visually inspected the genome-specific LPI histogram output by the algorithm (not-shown). A clear break point in the distribution of LPI frequencies was observed around value 0.5, which was then used as cut-off value, yielding 4655 LGT candidates.

A final examination of these candidates showed that there were many hits to sequence repeats. Accordingly, to remove this bias, cases where the number of different non-self BLAST hits was higher than 500 were removed from the candidate list, leaving 1092 prokaryotic and 2978 eukaryotic LGT candidates. Since the purpose of the present analysis was to evaluate the inter Domain lateral gene transfer to *Piromyces sp*. E2, only the Bacteria and Archaea candidates were further studied.

### Phylogenetic validation

Phylogenetic validation is the bona fide method to verify positive lateral gene transfer (LGT) candidates. This allows us not only to flag phylogenetically atypical genes, but it also provides information regarding their likely origin. Accordingly, we have computed Neighbor-Joining (NJ) trees for each of the 1092 prokaryotic LGT candidates (Figure 5).

To do so, for each candidate protein we have retrieved homologous sequences, using BLASTP (v2.2.25 default parameters), from two databases: NCBI’s nr database restricted to eukaryotic sequences, and UniProt’s uniref90 database, where each entry contains a sequence that represents a cluster of proteins with at least 90% sequence identity (to lower the amount of very similar hits, particularly for prokaryotic sequences). This approach was undertaken to ensure that our dataset contained homologous sequences from all domains (i.e. Eukaryota, Bacteria and Archaea) and not only prokaryotic hits due to the current bias towards these sequences in the sequence databases. Both BLASTP results were merged, and redundant sequences were filtered out. The sequences were subsequently aligned with ClustalW (v2.0.10) [74] and 100 times bootstrapped NJ trees computed with QuickTree (v1.1) [75].

To automatically process the 1092 trees we have devised a tree parsing algorithm (written in PERL and available upon request) that confirms the exclusive presence of prokaryotes in the second smallest partition containing Piromyces’ sequence, assuring that the target candidate sequence clusters within an exclusively “prokaryotic branch”. Because these are unrooted trees, this method assumes that the root can be anywhere, except between the smallest and the second smallest partition containing Piromyces’ leaf (method further detailed in [25]). From the initial 1092 trees, 704 passed this phylogenetic validation, and were further visually and individually inspected using FigTree (v1.3.1)(http://tree.bio.ed.ac.uk/software/figtree/)147 cl:779.

### LGT candidate annotation: Functional category assignment to orthologous groups

The final 704 lateral gene transfer candidates were assigned to one of the orthologous groups (COGs, KOGs or NOGs) from the STRING 8.0 database [76] using a BLAST sequence similarity search, applying a cognitor-like rule [77], as implemented in the Signature web server [78]. 577 out of 704 proteins were confidently assigned to at least one orthologous group (OG), totaling 207 different OGs (Supplementary data 2). Additionally, 395 of our candidates were present in at least one of the annotation records generated by the Piromyces’ genome sequencing team at the JGI: 358 had GO annotations, 85 had KEGG data and 183 had been assigned to KOGs (euKaryotic Orthologous Group) (Figure 5 and Supplementary data 2).

We pooled together all these data in order to obtain as many annotations as possible, and mapped the final dataset into NCBI’s COG function categories (“one-letter” code). KOGs and Signature OGs were directly assigned to one, or more, functional categories. GO accessions and KEGG Enzyme Commission (EC) numbers were first mapped to KOs (KEGG Orthology groups) and then, these KOs to COGs, which could then be assigned to a functional category. We have further manually classified the proteins that had at least one GO/KEGG annotation for which no automated functional category could be assigned. Finally, proteins without any source of annotation were classified as “S” – Poorly Characterized: Function Unknown. In the end, all 704 LGT candidate proteins had been assigned to at least one functional class, and there were 250 different OG annotations represented in the dataset.

#### LGT candidates: Pathway visualization and Functional category distribution

Using iPATH2 [61] we displayed the function relationships between the lateral transfer candidate genes against the background of *Piromyces sp*. E2 metabolic pathways. For Piromyces’ background metabolism, we used the list of the whole genome OG annotations (available at the Piromyces’ genome website), consisting of 8626 functionally annotated proteins successfully mapped into 2020 unique KOs, representing 868 metabolic edges. The 704 LGT proteins map to 201 unique KOs, defining 136 KEGG metabolic edges.

#### LGT candidates: Density as a measure of pathway cohesion

In order to find if the laterally transferred genes are metabolically coupled, we decided to assess the cohesion of the LGT metabolic graph and compare it to a set of randomly generated graphs sampled from KEGG’s Pan-metabolic map – the general metabolism map, containing all KEGG metabolic pathways. To this end, we performed a pathway cohesiveness analysis based on the graph density (d), calculated as follows:

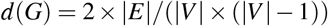

where *G* =(*E, V*) is an undirected graph with |*E*| number of edges (enzymes) and |*V*| number of vertices (metabolites). This measure is a good proxy for a graph’s group cohesiveness because it is a continuous metric bounded between zero and one, allowing the comparison between the values obtained for different sized graphs, while it also closely mirrors the visually perceived ‘‘pathway cohesion” (Figure 6).

**Figure 6.**
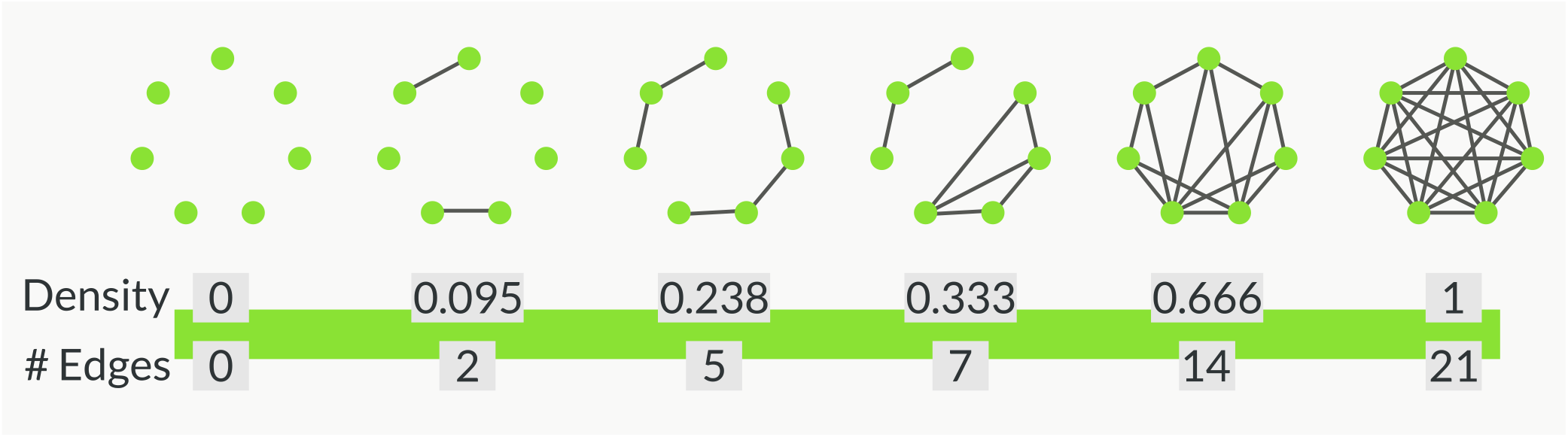
Graph density as a proxy for pathway cohesiveness. Diagram showing the visual correspondence between the graph density value and its overall cohesiveness, using a model graph with 7 nodes.

First, to calculate the observed-graph densities, we transformed the three pathway sets: LGT, Piromyces metabolism, and Pan-metabolism into undirected graphs where the edges are OGs and the nodes are metabolic compounds. Next, to evaluate the significance of the observed densities, we calculated the density distribution from 1 million random graphs composed by the sampling (without replacement) of N edges from a particular background graph. Three density distributions were generated: (i) 136 edges (the size of the LGT set) sampled from the Pan-metabolic graph; (ii) 136 edges sampled from the Piromyces’ background metabolism graph; and (iii) 868 edges (the size of Piromyces’ background metabolism) sampled from the Pan-metabolism map. (The R notebook with the annotated analysis pipeline is available online at https://doi.org/10.6084/m9.figshare.7173173.v2).

#### Assessment of LGT donor taxa

To confidently analyze the potential donor taxa, we selected the subset of highly supported candidates for which the second smallest partition branch presented a bootstrap value higher than 50% in the NJ tree. For the selected 240 proteins, Maximum Likelihood (ML) phylogenies were computed with RAxML (v7.2.8) [79], using the LG Substitution Matrix, a GAMMA model of rate heterogeneity with ML estimated alpha and 4 discrete-rate categories. The trees were 100 times bootstrapped, and 20 ML searches were conducted to obtain the best maximum likelihood tree.

All ML trees were computationally evaluated to find the last common ancestor for all non-Piromyces organisms present in the second smallest partition. The results were mapped into NCBI’s taxonomy tree to highlight the branches that contributed to our candidate LGT set. (All phylogenies computed for LGT assessment are available online as supplementary data 3: https://doi.org/10.6084/m9.figshare.7173020.v1).

## Acknowledgements (not compulsory)

This research was supported by (…)

ID was funded by the Portuguese Fundacao para a Ciencia e Tecnologia (SFRH/BD/32959/2006).

ID would like to thank REMM for stimulating scientific discussions and priceless scientific support.

## Author contributions statement

M.A.H. conceived the study, I.D. conducted the bioinformatics analyses, M.A.H. and I.D. analysed the results, I.D. and M.A.H. wrote the manuscript. All authors reviewed and approved the final manuscript.

## Additional information

## Supplementary information

All 4 supplementary data files have been deposited in figshare: https://doi.org/10.6084/m9.figshare.7173020v1.

Supplementary data 1: LGT candidate set: Hydrogenosomal protein prediction.

Supplementary data 2: LGT candidate set: Mapping to orthologous groups.

Supplementary data 3: LGT assessment: Phylogenies computed (in Newick format).

Supplementary figure 1: Distributions of the observed and random graph densities calculated.

## Competing financial interests

The authors declare no competing financial interests.

